# Genomic regions of durum wheat involved in water productivity

**DOI:** 10.1101/2023.06.07.544022

**Authors:** Meryem Zaïm, Zakaria Kehel, Miguel Sanchez-Garcia, Bouchra Belkadi, Abdelkarim Filali-Maltouf, Ayed Al Abdallat, Filippo Maria Bassi

## Abstract

Durum wheat is a staple food of the Mediterranean Basin, mostly cultivated under rainfed conditions. As such, the crop is often exposed to moisture stress. Therefore, the identification of genetic factors controlling the capacity of genotypes to convert moisture into grain yield (i.e. water productivity) is quintessential to stabilize production despite climatic variations. A global panel of 384 accessions was tested across eighteen Mediterranean environments (Morocco, Lebanon, and Jordan) representing a vast range of moisture levels. The accessions were assigned to water responsiveness classes, with genotypes ‘Responsive to Low Moisture’ reaching an average + 1.5 kg ha^-1^ mm^-1^ advantage. Genome wide association studies (GWAS) revealed that six loci explained the majority of this variation. A second validation panel tested under moisture stress confirmed that carrying the positive allele at three loci on chromosomes 1B, 2A and 7B generated an average water productivity gain of + 2.2 kg ha^-1^ mm^-1^. Interestingly, loci on chromosome 2A is novel. The three loci were tagged by Kompetitive Allele Specific PCR (KASP) markers, and these were used to screen a third independent validation panel composed by elites tested across moisture stressed sites. The three KASP combined predicted up to 34% of the variation for grain yield at 65% accuracy. These loci are now ready for molecular pyramiding and transfer across cultivars to improve the moisture conversion of durum wheat.

**Highlight:** Loci controlling drought tolerance were identified using a solid strategy, involving 3 different panels. Those loci associated enables higher water productivity and grain yield.

## Introduction

Durum wheat (2*n* = 28, AABB, *Triticum turgidum* L. ssp. *durum*) is a staple and cash crop grown on over 17 million ha worldwide (Tidiane *et al*., 2019; Xynias *et al*., 2020). Approximately 2/3 of durum wheat is grown in the Mediterranean basin, but this area contributes to only half of the worldwide production (Li *et al*., 2013; Kabbaj *et al*., 2017). In fact, climate change has and will continue to affect this region, with annual precipitation projected to decrease by 20–40% by the second half of the 21^st^ century (Zittis *et al*., 2021). Rainfall and temperatures in the Mediterranean dryland areas are largely unpredictable within and between cropping seasons. In the past years, North African countries have witnessed a raise in the frequency of drought events, an extension in their length, and an anticipation in their time of occurrence, substantially shifting from late spring to the middle of winter (Belaid *et al*., 2005; Tramblay *et al*., 2020; Qi *et al*., 2022). Since drought stress has a devastating effect on yield and its related traits (KiliÇ and Tacettin, 2010; Bilal *et al*., 2015), the North African durum wheat farmers have experienced strong reductions in their productivities. Under such conditions, breeders have committed to the delivery of new varieties with enhanced adaptation mechanisms, rather by avoiding or tolerating these stresses. Genetic improvement programs have for a long time attempted to balance the needs of raising overall yield potential, while ensuring cultivars with stable yield performances across seasons. In fact, the final productivity of a variety results from the combined effects of genotype (G), environment (E), and their interaction (GxE) (Mohammadi *et al*., 2015). Thus, the development of superior cultivars requires strategic approaches to combine good stress tolerance with strong yield stability (Mohammadi *et al*., 2011; Bassi and Sanchez-Garcia, 2017).

Yield stability refers to the ability of certain genotype to ensure good yield performances despite the fluctuations of growing conditions occurring across environments, and it is normally associated with the GxE component. Several decades of studies have demonstrated that stability is controlled by genetic factors interacting with the environment. As such, it is possible to improve the stability of a genotype via pyramiding multiple positive alleles for this trait. Breeders approach this need by testing the genotypes under a vast range of environments and seasons, to then derive what are defined as stability scores (Malosetti *et al*., 2013) and then use these to identify stable genotypes across environments. One such score widely used in durum wheat breeding is the AMMI wide adaptation index (AWAI) score that utilizes the AMMI capacity to partition the GxE into sub-factors, to then estimate a weighted value to be assigned to the genotype (Bassi and Sanchez-Garcia, 2017). However, a stable variety can also be obtained by pyramiding multiple positive alleles at loci controlling discrete interactions with the environment. For instance, a drought tolerant variety would be able to maintain its yield performance (i.e. stability) even when moisture stress occurs. The concept of water productivity is linked to yield stability and potential as it has been used in plant breeding to define genotypes capable of using moisture in a more efficient way, and hence achieve higher productivity at the same level of moisture input (Anyia *et al*., 2008). While the application of this concept was originally proposed to define the response of genotypes to increasing irrigation rates, it has become even more important to assess the response to moisture stress, when water availability is particularly scarce (Bhouri Khila *et al*., 2021). In fact, wheat’s most sensitive growth stages to water stress are mainly stem elongation and booting, followed by anthesis and grain filling (Blum and Pnuel, 1990; Shpiler and Blum, 1990; del Moral *et al*., 2003). Water deficit around anthesis may lead to a loss in yield by reducing spike and spikelet number and the fertility of surviving spikelets, while water deficit during grain-filling period reduces grain weight (Karam *et al*., 2009). Geerts and Raes (2009) indicate that scarce moisture can increase water productivity for various crops without causing severe yield reductions. Zhang *et al*. (2006) demonstrated that under rainfed conditions, wheat grain yield, harvest index and water productivity were greatly improved under regulated deficit irrigation when compared to the non-water stressed treatment. Maximizing WP may be economically more profitable for the farmer than maximizing yields or land productivity (English, 1990) in areas where water is the most limiting factor. Karrou and Oweis (2012) results showed that in general a reduced irrigation of 1/3 of full supplemental irrigation gave the highest rate of increase in grain yield and water productivity. Grain yield reductions due to the application of 2/3 supplemental irrigation were around 10% of the full supplemental irrigation on average. While differences in total water productivity of crops grown under full irrigation compared to deficit irrigation were not significant.

Beyond the application of stability systems to adapt to all conditions, there are several discreet traits that have been proposed as favoring the adaptation of durum wheat to moisture stress. A simplified list of these would include early maturity to avoid terminal stress (Gupta *et al*., 2020), good coverage of ground to favor shading and prevent moisture transpiration from the soil (Yadvinder *et al*., 2014), access residual moisture in deeper soil layers (Yadvinder *et al*., 2014; Lilley and Kirkegaard, 2016; El Hassouni *et al*., 2019), and improvement of specific yield components, with a particular attention to grain size (Mohammadi *et al*., 2019). In that sense, breeders seek to identify and pyramid these traits to achieve better stability when moisture stress occurs (Araus *et al*., 2008; Reynolds *et al*., 2009; Tuberosa, 2012; Sukumaran *et al*., 2018). Therefore, the knowledge of genetics and gene action of these traits is essential for generating stable varieties (Habash *et al*., 2009). Molecular markers technology offers the possibility to identify and track these positive alleles (Collard and Mackill, 2008; Ceccarelli, 2015). Genome wide association study (GWAS) is an approach that helps determine significant relationships between the allelic make up (i.e. haplotypes) of a genotype and its field response. Such approach was used for the identification of novel QTLs with potential implications for durum wheat breeding programs, such as loci associated with variation in kernel size (Fiedler *et al*., 2017), grain yield and its components (Mangini *et al*., 2018; Sukumaran *et al*., 2018; Wang *et al*., 2019), but also response to moisture changes and roots. Maccaferri *et al*. (2011) used an association mapping to dissect the genetic basis of drought-adaptive traits and grain yield (GY) in a collection of 189 elite durum wheat accessions evaluated in 15 environments highly different for water availability during the crop cycle (from 146 to 711 mm). For GY, significant associations were mostly detected in one environment only, while decreasing rapidly from two to five environments and with only one marker found significant in six environments. While in another study, Maccaferri *et al*. (2016) used linkage and association mapping for root system architecture in two recombinant inbred line populations and one association mapping panel of 183 elite durum wheat accessions evaluated as seedlings revealed 20 clusters of QTLs for root length and number, as well as 30 QTLs for root growth angle (RGA). Divergent RGA phenotypes observed by seminal root screening were validated by root phenotyping of field-grown adult plants.

In the present study, we aimed at broadening the understanding of the genetic factors involved in controlling water productivity and moisture stress adaptation in durum wheat. Therefore, we investigated the performances of a large ‘discovery panel’ of durum wheat accessions across 18 environments experiencing different degrees of in season moisture. Beyond the identification of stable and top performing entries, this investigation sought to define discrete clusters of water productivity types. GWAS on the ‘discovery panel’ was then used to identify haplotypes more frequently present in the most water responsive genotypes, which were then investigated in a second ‘confirmation panel’. Finally, to ensure these critical loci can be readily incorporated into novel cultivars, Kompetitive Allele Specific PCR (KASP) markers were developed to tag them and then confirmed for their ability to predict moisture stress adaptation on a third ‘validation panel’.

## Material and methods

### Plant material

This study evaluated three discrete germplasm panels. The first is defined as the ‘discovery panel’ and it includes 384 durum wheat entries including landraces, elites and cultivars (Tab S1). The kinship of this panel was previously presented by Kabbaj *et al*. (2017), and it has already been used to identify the genomic loci involved in resistant to a damaging insect pest (Bassi *et al*., 2019), phenology (Gupta *et al*., 2020), and its response to heat stress (El Hassouni *et al*., 2019). This panel was tested in its entirety at some environments, while a subset was used in other environments as explained in more details below. The second set of entries is defined as the ‘confirmation panel’ and it includes 94 ICARDA’s elites that constituted the 2019 international nurseries 42^th^ International Durum Observatory Nursery (IDON; Tab S2). The third set is defined as the ‘validation panel’ and it includes 94 ICARDA’s elites that constituted the 2020 international nurseries 43^th^ IDON (Tab S3).

### Field trials and management

The ‘discovery panel’ was assessed during 2014-15, 2015-16, 2016-17 and 2017-18 growing seasons in eighteen contrasting environments as described in **Fig 1**. Four were located in Morocco: Marchouch (MCH), Sidi El Aidi (SAD), Melk Zhar (MKZ) and Tessaout (TES), two in Lebanon: Terbol (TER) and Kfardan (KFD) and one in Jordan: Musghar (MUS). The experimental design was an augmented design with four replicated checks in the 2014-15 (15) and 2015-16 (16) growing seasons in MCH15, MCH16, SAD16, MKZ15, MKZ16, TES16, TER15, TER16, KFD16, and MUS18 during 2017-18 season (18). During 2016-17 (17) and 2017-18 (18) seasons, a subset of 144 genotypes was selected and used to run an alpha-lattice design with two replications and twelve incomplete blocks of size twelve, at MCH17, MCH18, SAD17, TES17, KFD17 and KFD18. Each entry was planted in plots of 6 rows of 5 m in length, row spacing was 0.2 m, for a total sown surface of 6 m^2^ at a seeding rate of 120 kg ha^-1^. Agronomic practices generally follow a timely sowing date between 15^th^ of November to 15^th^ of December with a base pre-sowing fertilizer application of 50 kg ha^-1^ of N, P and K. Planting occurred after a legume crop season. During 2016-

**Figure 1.**
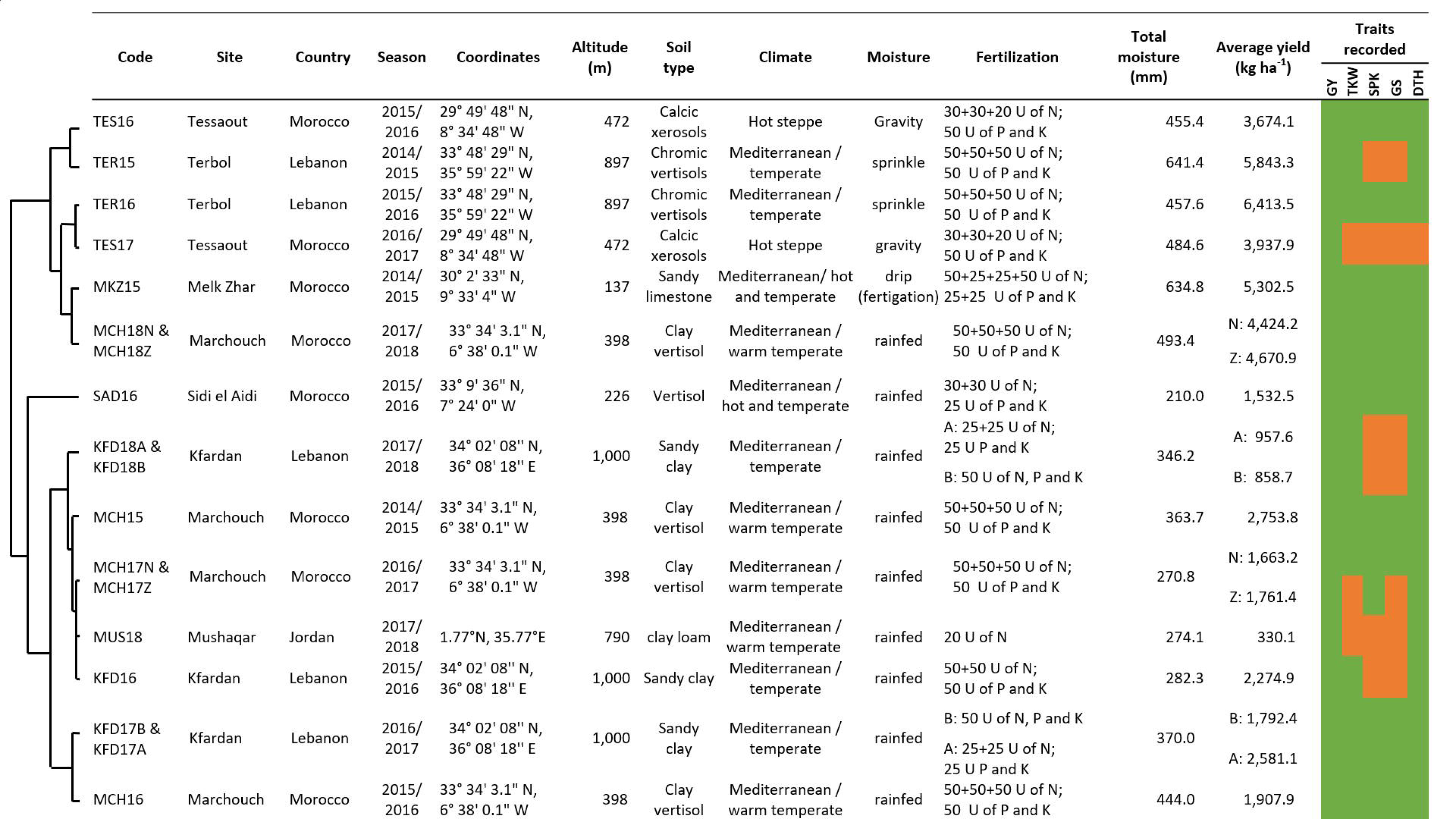
Description of the testing environments used for the ‘discovery panel’, and their principal component differentiation (PCA) hierarchical clustering based on climatic factors. GY: Grain yield (kg ha^-1^), TKW: 1,000 Kernel weight (g), SPK: Spike density per m^2^, GpS: Grain per spike, PLH: Plant height (cm), DTH: Days to heading.

17 and 2017-18 seasons in MCH, two management conditions have been used: normal sowing (MCHN) following standard land preparations and tillage, and zero tillage (MCHZ) on a fully retained faba bean stubble. Both sowings were conducted using the same seeder, even though it was specifically developed for zero till practices. At stage 14 of the Zadok’s scale (Z) herbicide was applied in a tank mixture to provide protection against both monocots and dicots. A week after herbicide application, ammonium nitrate was provided to add 36 kg ha^-1^ of N. When in season moisture exceeded 350 mm a final application of urea was used at flowering to deliver additional 46 Kg ha^-1^ of N. In KFD17 and KFD18, two kind of fertilizer applications were done: KFDA with only basal fertilization 50 kg ha^-1^ of N, P and K and KFDB with additional 50 kg ha^-1^ of Urea at Z15. In MKZ, the first basal fertilization was followed by 5 split applications each of 20 Kg ha^-1^ of N via fertigation through drip pipes. Three sites were irrigated: TES, where four gravity irrigations of 35 mm each were provided after Z10, Z18, Z45, and Z65; MKZ, where 12 irrigations of 10 mm each were provided via drip irrigation at one week interval from two weeks after Z10 to Z89 and TER, where two sprinkle supplemental irrigation of 20 mm each were provided before Z10 and after Z65. The remaining experiments were conducted under rainfed conditions with total rainfall values and other details presented in **Fig 1**.

The station of Sidi El Aidi (SAD) in Morocco was identified by the global initiative CRP WHEAT as an ideal site to test for drought tolerance of wheat, and for this reason it was also used to screen the two other panels. The ‘confirmation panel’ and the ‘validation panel’ were tested at SAD as augmented design with four replicated checks, during seasons 2018-19 and 2019-2020, respectively. The total moisture recorded during 2018-19 and 2019-2020 was 296 and 286 mm, respectively, which constitute strong moisture stress for durum wheat.

### Phenotyping

Days to heading (DTH) was recorded as days elapsing between sowing and 50% of plants showing emerging heads. At maturity, the number of fertile spikes were counted in 0.25 m^2^ and this value was multiplied by four to derive the number of spikes per m^2^ (SPK). Grain yield (GY, kg ha^-1^) was recorded by harvesting the central four rows of each plot, weighting it on a precision scale and dividing this value by the plot surface. From the harvest of each plot, 1,000-kernel weight (TKW, g) was determined by counting five hundred randomly selected grains on a ‘Choppin Numigral’ counter followed by weighting on a precision scale. The number of grains per meter square (Gr.m^-2^) was calculated using the total weight of the plot, divided by the harvested surface and the estimated weight of one kernel, derived from the TKW value, as per (1):

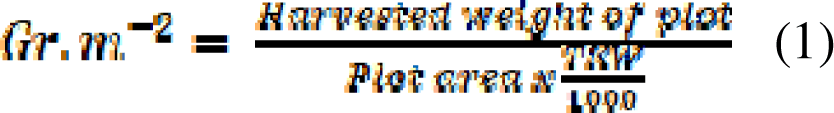

The number of grains per spike (GpS) was then derived by dividing the number of grains per meter square (1) by the number of spikes recorded for the same area as follows (2):

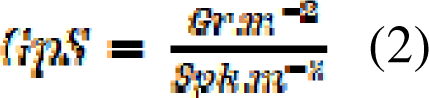

As detailed in **Fig 1**, GY was recorded in all environments, while the other traits were collected only in some.

### Field data analysis

Analysis of variance was performed using Genstat for the augmented designs, while alpha lattice and the combined analysis were run on GEA-R 4.1 in the R environment (Pacheco *et al*., 2015). Combined ANOVA across mega-environments was obtained by linear model fitted considering genotypes as fixed term (Team R. Core, 2017). Best linear unbiased estimates (BLUEs) were calculated for each genotype at each environment defining genotypes as fixed effect using the R package ASReml-R (Butler *et al*., 2009). The package ASReml-R was also used to estimate the narrow-sense heritability. Broad-sense heritability was calculated separately for each design by Genotype x Environment Analysis with R (GEA-R) version 4.1. The ratio of variance accounted by each source of variations (G, E, and GxE) was calculated dividing the sum of square of each source by the total sum of the square.

For grain yield, GxE was partitioned by additive main effects and multiplicative interaction (AMMI) model using R package Agricolae (De Mendiburu and Yaseen, 2020). The ‘AMMI wide adaptation index’ (AWAI) measures the distance of each genotype from each significant IPCs axis and it was calculated using the following formula, presented by Bassi and Sanchez-Garcia (2017):

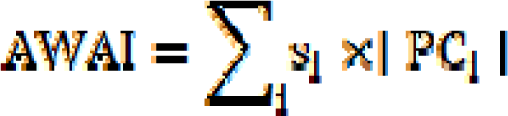

Where *i* is the number of significant IPCs determined by classical F-test in R (Team R. Core, 2017), *s_i_* is the percentage of total GxE variance explained by each IPC, and PC is the actual IPC value. AWAI values close to ‘0’ are obtained for the most widely adapted and stable germplasm (Malosetti *et al*., 2013). As indicated by Bassi and Sanchez-Garcia (2017), a biplot between the genetic (G) component of yield (i.e. yield potential) and the interaction (GxE) component (i.e. yield stability) was used to determine the best genotypes combining both G and GxE for grain yield. The AWAI index explaining GxE was presented as ratio to minimum value, and values close to ‘1’ were obtained for the most widely adapted and stable genotypes. To define the genetic component of GY obtained across environments with vast differences in the average performances, the actual values were converted to a ratio of the top performing entry at each environment and then averaged across.

A climate matrix was developed for each environment, splitting the records into five growth stages: one month before sowing, sowing until the end of the vegetative stage, flowering stage, grain filling period, and physiological maturity period. Simple linear regression was conducted between the climatic matrix and the response of genotypes at each site for GY. The climatic factors having a significant effect (p < 0.05) were used to perform hierarchical clustering among environments using the R package FactoMineR (Josse *et al*., 2008) (**Fig 2)**.

**Figure 2.**
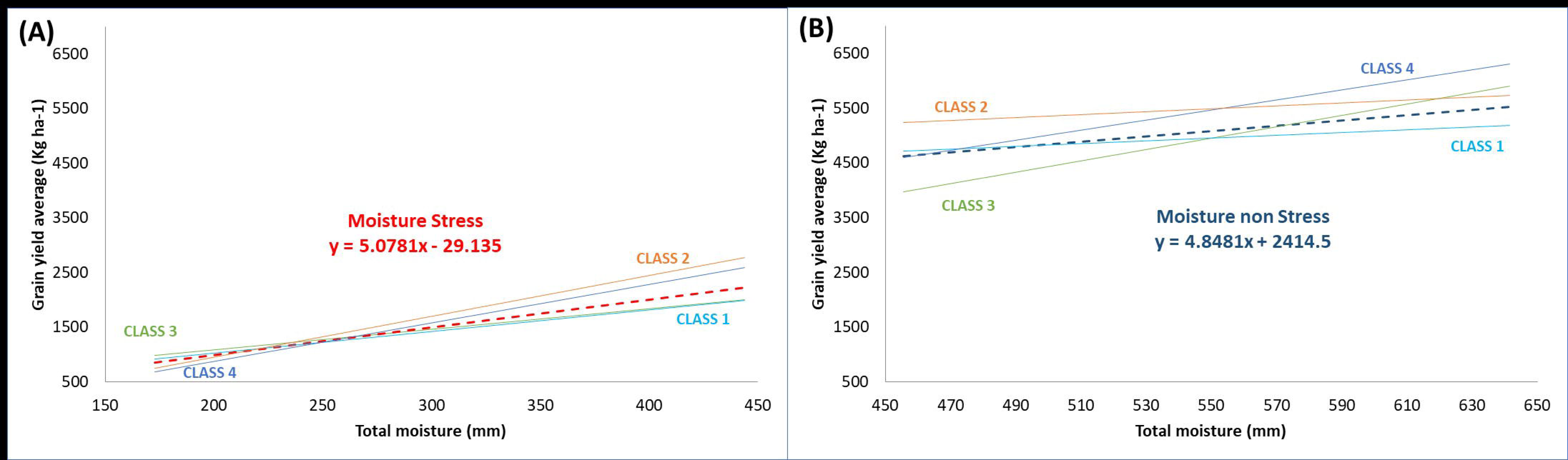
Trendlines of linear regression representing water productivity as biplot of the total moisture at each environment against the grain yield at that environment. One representative genotype example for each water productivity class (Tab 1) is presented and compared to the average performance of all genotypes at each environment (dashed line). Trendline color cyan represent the ‘class 1’, the orange for ‘class 2’, green for ‘class 3’ and blue for ‘class 4’, Moisture stressed environments are presented at (A) and non-stressed at (B).

### Assignment of genotypes to water productivity classes

The water productivity (WP) is calculated using the following formula:

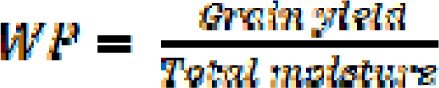

To define the average trend, the average GY performances of each environment was plotted against the total moisture of that environment, which substantially corresponds to a graphical representation of the average WP trend. To increase the accuracy of this trend, the environments were split into two groups, one defined as ‘stressed’ including 11 environments experiencing moisture stress, and the second defined as ‘non-stressed’ including seven environments where moisture stress did not occur. The slope (b) of WP was calculated for each group, reaching 5.08 for moisture stress cluster and 4.85 for non-moisture stress. These values represent then the hypothetical average performance of a given genotype tested at that group of clusters. Hence, higher values (steeper response to water increase) would be obtained by genotypes with higher WP, while lower values (flatter curve) would be associated to less responsive genotypes.

To assess this, the GY performance of each genotype at each environment was plotted against the moisture level of that environment and the actual slope value (b_i_) for the two trend lines (moisture stressed and non-stressed) were calculated (**Fig 2**). Based on these b_i_ values, genotypes were assigned to four different water responsive classes representing more or less responsiveness compared to the average trend (Table 1). However, to ensure that the trendline explained the observed changes in moisture, genotypes for which the regression value between GY and moisture levels was not significant (p<0.01) were assigned to a fifth class of water unresponsive behavior.

**Table 1.**
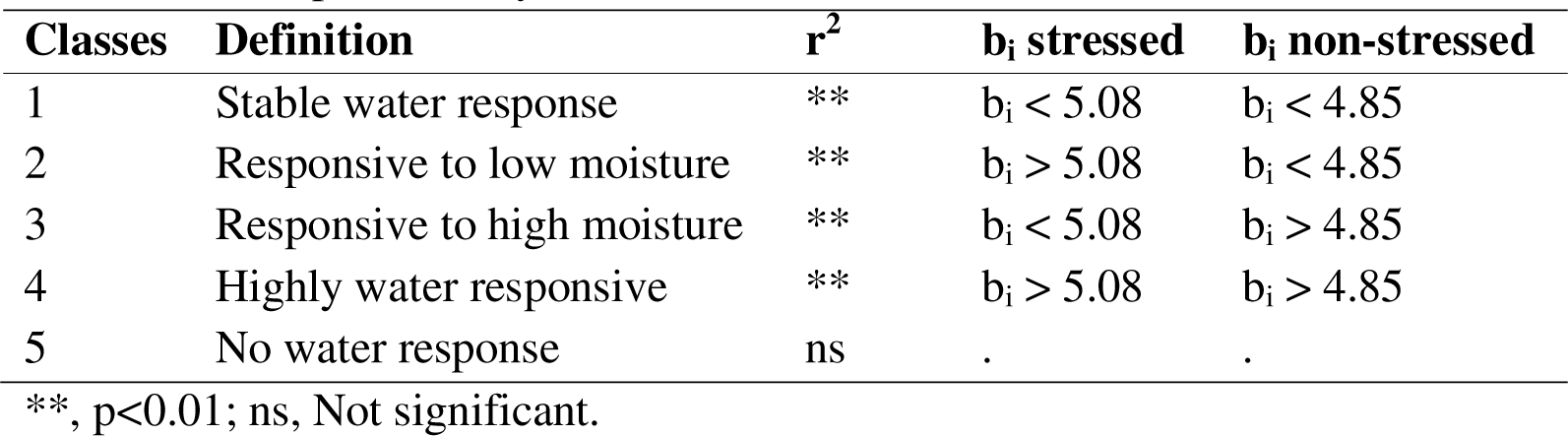
Water productivity classes

| **Classes** | **Definition** | **r<sup>2</sup>** | **b<sub>i</sub> stressed** | **b<sub>i</sub> non-stressed** |
| --- | --- | --- | --- | --- |
| 1 | Stable water response | ** | b<sub>i</sub> < 5.08 | b<sub>i</sub> < 4.85 |
| 2 | Responsive to low moisture | ** | b<sub>i</sub> > 5.08 | b<sub>i</sub> < 4.85 |
| 3 | Responsive to high moisture | ** | b<sub>i</sub> < 5.08 | b<sub>i</sub> > 4.85 |
| 4 | Highly water responsive | ** | b<sub>i</sub> > 5.08 | b<sub>i</sub> > 4.85 |
| 5 | No water response | ns | . | . |

\*\*, p<0.01; ns, Not significant.

### Genotyping and association mapping analysis

The ‘discovery panel’ was genotyped with 35K Affymetrix Axiom wheat breeders array (www.affymetrix.com) to generate 7,652 polymorphic SNP with 98 to 100% identity when blast aligned to the Svevo genome (Maccaferri *et al*., 2019), less than 1% missing data, minor allele frequency higher than 5% and heterozygosity less than 5% as detailed in Kabbaj *et al*. (2017). These authors also defined a kinship structure of 10 sub-clusters. Genome wide linkage disequilibrium (LD) decay analysis were performed by Bassi *et al*. (2019) and defined as 51.3 Mbp. GWAS was performed for the panel using as phenotypic input the BLUEs of each trait at each environment, and using the BLUEs estimated from the combined analysis of the two moisture groups (stressed and non-stressed). TASSEL 5 software (Bradbury *et al*., 2007) was used for the analysis imposing DTH as covariate to avoid identifying flowering genes, since these were already described in Gupta *et al*. (2020). Two models were used and compared using two additional covariate parameters, Q (population structure) and K (Kinship). Q model was performed using a general linear model (GLM), and Q + K model using a mixed linear model (MLM). The best model for each trait was selected based on the quantile-quantile (Q-Q) plots (Sukumaran *et al*., 2012). Significant marker-trait associations (MTA) were determined using a Boneforroni correction by LD as suggested by Duggal and Beaty (2008) for p < 0.05 corresponding respectively to LOD = 2.69 (Bassi *et al*., 2019). In addition, Pearson’s critical values (Pearson, 1895) for correlations r was squared to obtain a critical r^2^ of 0.024 (p<0.01) and used to determine significant markers explaining sufficient ratio of the total phenotypic variation. Any marker-traits associations (MTAs) with LOD and r^2^ superior to these cut-offs were considered valid and presented here. MTAs falling at a distance inferior to twice the LD (102.6 Mbp) were deemed to be too physically close to be resolved by this panel into distinct loci and hence were assigned the same QTL identifier. Consistent QTL gather MTA appeared both in the combined and more than one individual environment.

The ‘confirmation panel’ was genotyped using a 23K array chip developed by SGS - Institut Fresenius TraitGenetics Section (Germany) which incorporates 14.5K SNPs from the 90K Infinium Array, 8.5K SNPs from the Axiom array, and 265 SNPs reported as linked to genes in the literature (Vitale *et al*., 2021). Marker curation was conducted as for the ‘discovery panel’, to result in 6325 polymorphic SNPs. These were also aligned to the Svevo genome assembly (Maccaferri *et al*., 2019). A kinship structure of 8 sub-clusters was identified (Table S2) and linkage analysis revealed that the LD was 21.2 Mbp, which resulted in a significant r^2^ = 0.05 (p<0.01). The two genotyping platforms were merged using the Svevo genome assembly as scaffold. Similarly to ‘discovery panel’, GWAS was conducted using flowering time as covariate. Significant MTA were determined using Boneforroni correction for p < 0.05 corresponding to LOD = 4.1. ShinyCircos software (Yu *et al*., 2018) was used to graphically represent the MTA and QTL identified by both ‘discovery’ and ‘confirmation’ panels.

The most representative marker for each QTL identified by the ‘discovery panel’ was selected based on its higher LOD and explaining a broader fraction of the phenotypic variation. Discrete classes of genotypes from the ‘confirmation panel’ were then defined based on their allelic combinations at the representative markers. These classes were defined as ‘haplotypes’. The phenotypic performances of the ‘confirmation panel’ genotypes belonging to each haplotype class were defined as a random effect, and a linear model was run to determine significance difference by LSD using LSD.test function of *agricolae* package (Team R. Core, 2017; De Mendiburu and Yaseen, 2020).

The 35K and 25K array probe sequences underlying the most interesting QTLs were submitted to LGC to run their proprietary software to assess their suitability to design KASP primers. For each QTL, four potential primer sets were synthetized and run on the ‘validation panel’. For each KASP marker that amplified and showed polymorphism, its allelic score was regressed against the GY value and a significance threshold was set at r^2^ >0.105 (p<0.01). In addition, the top 20 yielding genotypes were defined as the ‘positive’ cases and the worst 20 genotypes as the ‘negative’ cases. The marker score was then evaluated among the positive and negative cases to define correct SNP call (true positive or true negative) and the wrong SNP calls (false positive and false negative). The marker accuracy was then calculated as the ratio of the correct allelic call among all, sensitivity as the ratio of the correct positive allelic calls among all, and specificity as the ratio of the correct negative allelic calls among all. The primer sequence of the markers is protected by commercial rights and cannot be disclosed here, but these can be purchased by all users as service via LGC indicating the marker names provided here.

## Results

### Phenotypic variation under moisture stressed and non-stressed environments

Analysis of variance revealed significant differences (p < 0.01) for the genotypes (G), environments (E), and their interaction (G×E) for most of the traits (Tab S4). The E effect explained most of the variation for GY (73%), GpS (69.3%), TKW (83%), SPK (88%) and DTH (84%), while the G effect explained the largest variation for GpS (16%). The G×E interaction showed a larger contribution to the total variability compared to the G effect for GY and SPK. Good heritability was obtained at all environments for all traits.

Vast phenotypic variation was recorded for all traits across the eighteen environments (Fig S2). The highest average GY was recorded in TER16 (6,413.5 kg ha^-1^), while MUS18 had the lowest average GY (330.1 kg ha^-1^) (Fig 1). Moisture data shows patterns of variation across environments, with some sites having a prevalence of drought events (MUS, KFD, SAD, MCH, Fig 1). GY performances were significantly (p < 0.05) influenced by the total water input during vegetative, flowering and grain filling stages and the maximum temperature during the flowering stage (Tab S5). Because of their significant effect on yield, these climatic factors were used to cluster the environments by PCA in two mega-environments: *i.* moisture stressed (MUS18, KFD17, SAD16, MCH15, MCH17, KFD16, MCH16 and KFD18) and *ii.* non-moisture stressed (TES16, TER15, MKZ15, TES17, TER16 and MCH18).

To avoid range effects, grain yield (BLUE) was converted to ‘ratio to the max’, to scale the variation based on the best performing entry. Under moisture stressed conditions the CIMMYT line DURUM_PANEL_UNIBO-016 (GID: 800032262, 3040.6 kg.ha^-1^) was the top yielding. Among the top highly performing entries, the ICARDA wide crosses DWAyT-0306 (GID: 800032191, 2563.5 kg.ha^-1^), DAWRyT-0106 (GID: 800043267, 2375 kg.ha^-1^), Magrour (GID: 800032178, 2241.6 kg.ha^-1^) and the elite lines MCHCB-0133 (800032351, 2204 kg.ha^-1^) and Icakassem1 (GID: 800030179, 2191.4 kg.ha^-1^). The top yielding line under non moisture stressed conditions was the ICRADA elite IDON37-094 (GID: 800043103, 7516.9 kg.ha^-1^), the Moroccan line Ourgh (GID: 4984522, 6407.1 kg.ha^-1^) was also among the highly yielding.

Partitioning the GxE effect by AMMI defined seventeen significant components (PCs), of which the first three combined accounted for 67.9% of the variation. The definition of the AWAI score determined an average performance equal to 0.2, with the two most stable lines being DURUM_PANEL_UNIBO-012 (GID: 800032258) a CIMMYT elite line and MCHCB-0133 (GID: 800032351) an ICARDA elite line. The bi-plot combining GY performances across sites and stability (AWAI) provides an ideal selection index to combine G and GxE effects (Fig 3) (Bassi and Sanchez-Garcia, 2017). Combined analysis under moisture stress identified 24% of genotypes having higher than GY and AWAI average. DURUM_PANEL_UNIBO-016 (GID: 800032262), DURUM_PANEL_UNIBO-017 (GID: 800032261) and AMEDAKUL1 (GID: 800032280) were the top3 stable and yielding. The ICARDA elite line Icakassem1 (GID: 800030179) and DWAyT-0306 (GID: 800032191) were also among the highly performing. While under non-stress conditions, 32% of the tested entries had higher yield and AWAI than the average. The highly yielding genotypes IDON37-094 (GID: 800043103) and Ourgh (GID: 4984522) were not stable. Contrary to the australian elite line Jandaroi (GID: 800032336), CIMMYT line DURUM_PANEL_UNIBO-024 (GID: 800032267) and the ICARDA line MCHCB-082 (GID: 800032342) were the top 3 stable and highly performing.

**Figure 3:**
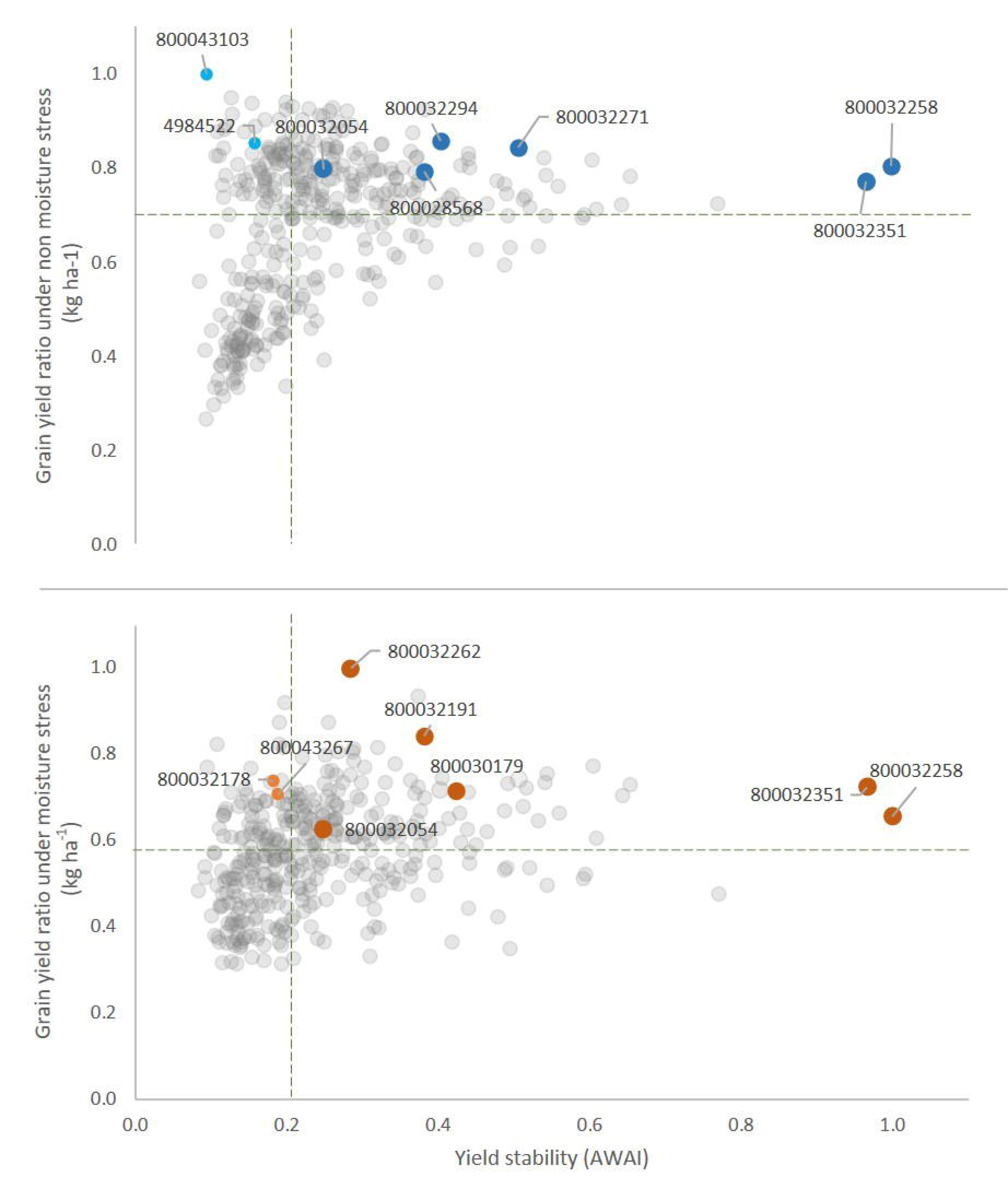
AMMI wide adaptation index (AWAI) against the ratio to the max of yield potential across 11 moistures stressed (up) and 7 no-stressed (down) environments. Dashed lines trace the average for each axis.

Beside stability per se, several traits contribute to the adaptation of genotypes to the environment. To determine which traits contributed to GY variation, correlation analysis was performed for each individual and mega-environment. This interaction (Table 2) revealed that GpS influenced (p < 0.001) GY in all environments. TKW had an effect only in 5 out of 9 moisture stressed environments and in all the non-stressed ones. Overall, SPK was not significantly correlated with GY under non moisture stress, only two environments showed highly significant relationship, while it accounted for 60% of yield variation in moisture stressed environments. In most of the environments flowering time shows a highly significant correlation with GY.

**Table 2:**
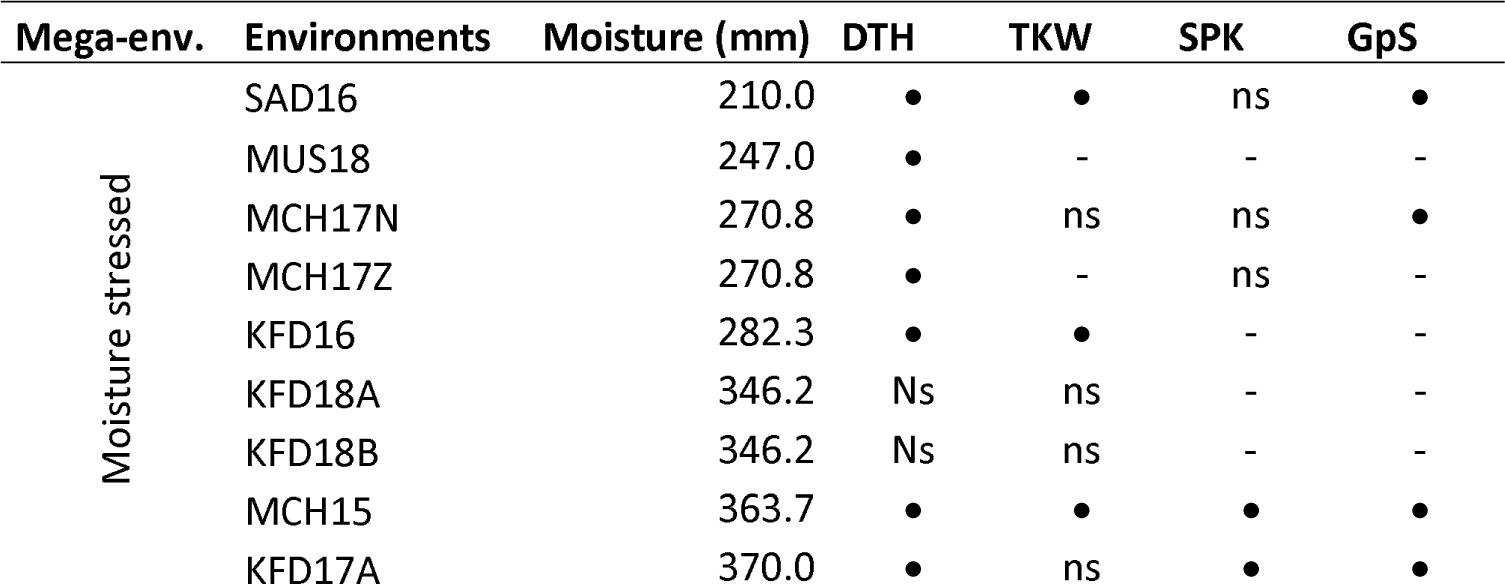
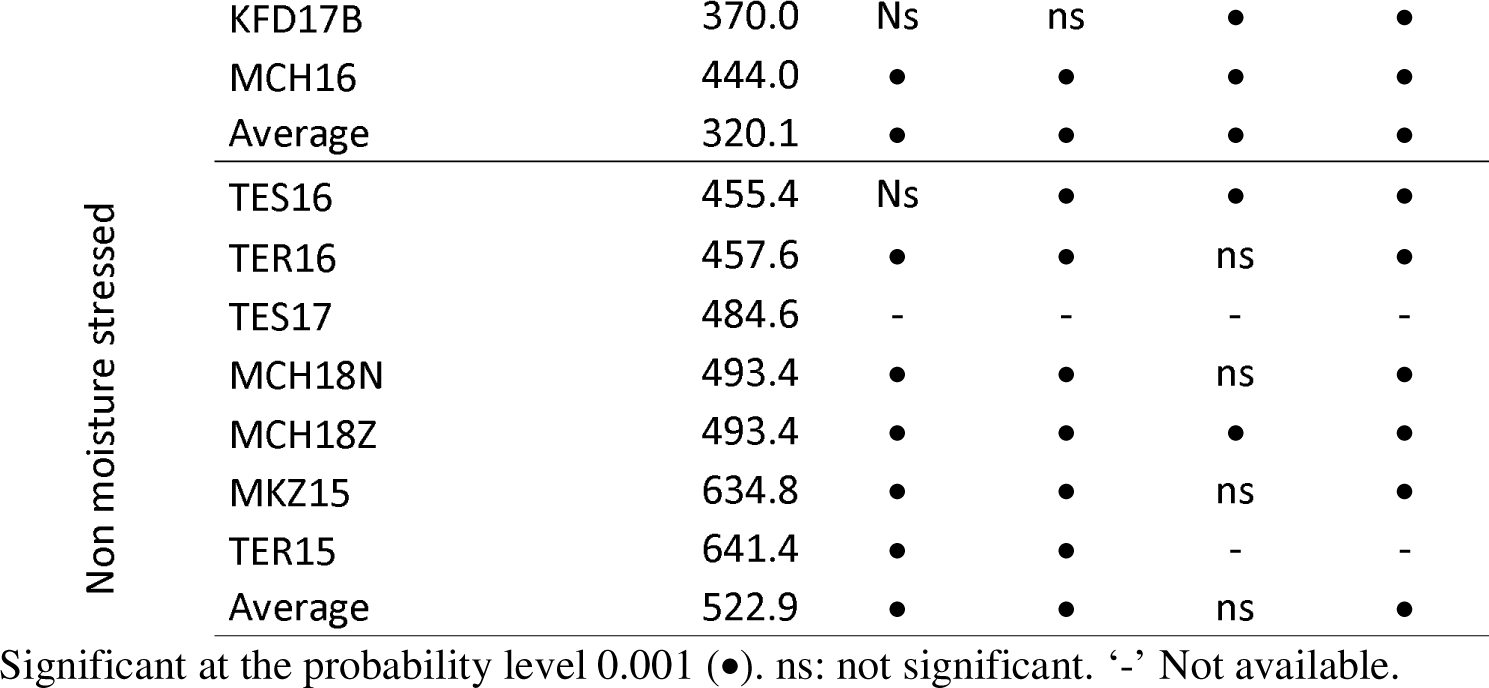
Correlation analysis for all traits against grain yield across moisture and no-moisture stressed conditions.

### Water productivity performance of genotypes

Climatic regression against GY identified that moisture amount during the vegetative, flowering and grain filling stages are the most significant climatic factors, explaining more than 70% of its variation (Tab S5). To better elucidate the relationship between GY and moisture, a water productivity (WP) value was calculated for each genotype. The average WP was estimated at 7.2 kg ha^-1^ mm^-1^ showing a significant linear relationship (r^2^ = 0.327) to the increase in moisture levels, resulting to an average WP of 5.1 kg ha^-1^ mm^-1^ across moisture stressed environments, while it reached 9.7 kg ha^-1^ mm^-1^ across non stressed environments (Fig S3).

A subset of 120 genotypes that have been assessed at all environments, was assigned to WP classes based on their respective trend of yield variations plotted against the moisture levels across environments. One quarter of the tested entries were assigned to class 3 ‘Responsive to high moisture’ and one quarter to class 4 ‘Highly water responsive’, while class 2 ‘Responsive to low moisture’ incorporated 20% of genotypes, 18% belong to class 5 ‘No water response’ and 13% as class 1 ‘Stable water response’. From a breeding perspective, class 2, 3, and 4 are the most interesting because they identify genotypes capable of producing more yield per water input compared to the average. Interestingly, Icakassem1 (GID: 800030179), DAWRyT-0106 (GID: 800043267) and Magrour (GID: 800032178) resulted among the highly WP performing elite lines under moisture stress, belonging to class 2 ‘Responsive to low moisture’. While under non moisture stress, the ICARDA elite line Miki3 (GID: 800028568) belonging to class 3 ‘Responsive to high moisture’ was among the highest for WP. Instead, DURUM_PANEL_UNIBO-023 (GID: 800032054), a CIMMYT line with top yield under both moisture conditions, belongs to class 4 ‘Highly water responsive’ (Table S6).

### QTLs controlling traits under moisture stressed and non-stressed conditions

A total of 280 significant MTAs were identified across individual and cluster of environments for all tested traits. The MTAs explained from 3% to 22% of the phenotypic variation and LOD ranged from 2.7 to 7.2. MTAs were distributed across 47 discrete QTL (Fig 4, Tab S7).

**Figure 4.**
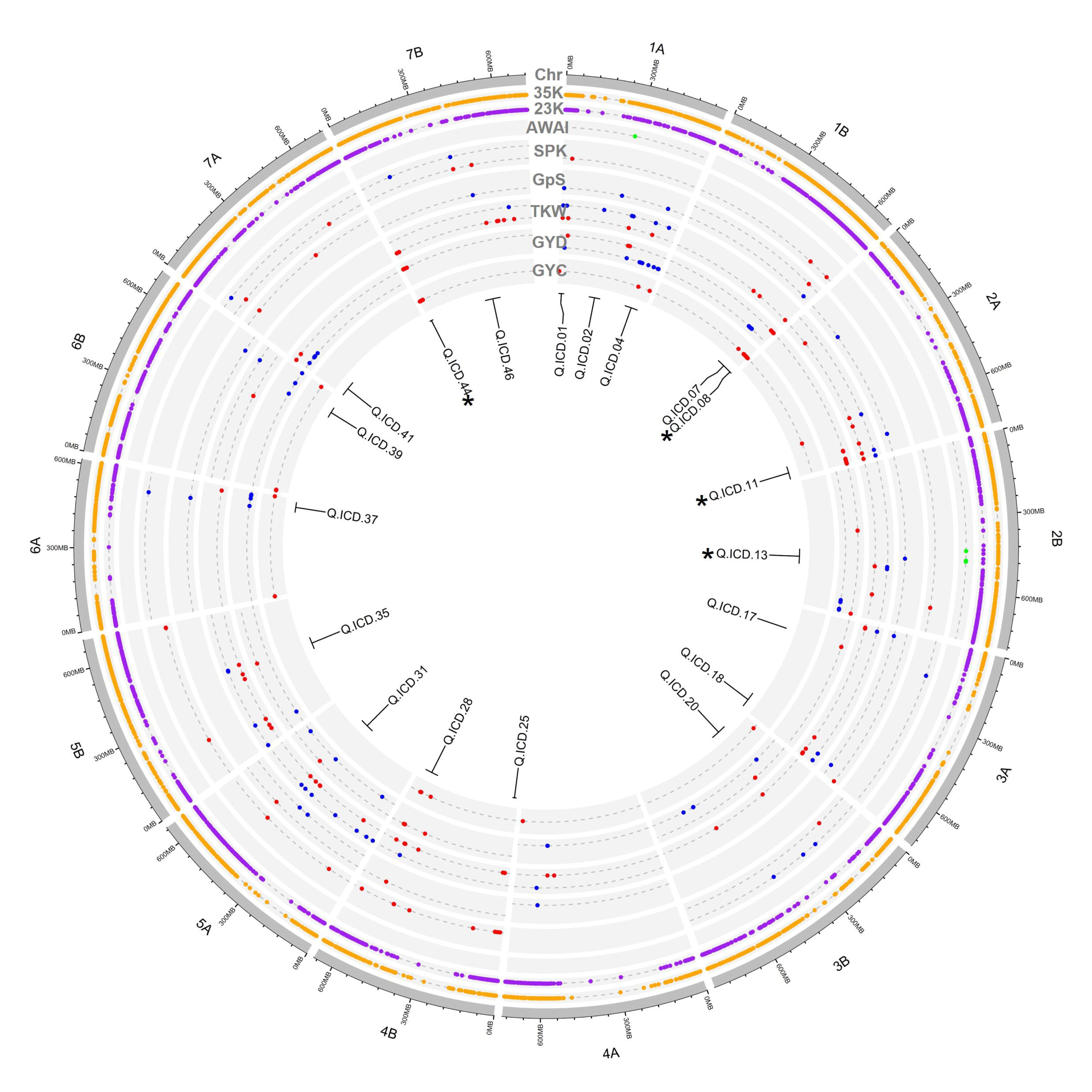
QTL discovery. The outermost circle shows the Svevo durum wheat genome assembly (Macaferri et al. 2019), including its chromosomes, followed by the distribution of 35K Axiom polymorphic probes in the ‘discovery panel’ (35K) and the 23K SNP probes in the ‘confirmation panel’ (23K). The following tracks represent results of significant marker trait association (MTA) identified in the ‘discovery panel’ for AMMI wide adaptation index (AWAI), spike density per m^2^ (SPK) and grain per spike (GpS), 1,000 kernel weight (TKW) and grain yield (GYD) across individual and combined environments. The following tracks represent MTA for GY identified in the ‘confirmation panel’ (GYC). The MTA identified in moisture stressed environments are color coded as red dots, while those identified in non-moisture stressed conditions are coded as blue dots. The innermost circle provides the QTL labels for reference. * represents the QTL confirmed by the GYC.

Under non-moisture stress environments, four consistent QTLs (Q.ICD.04, Q.ICD.07, Q.ICD.37 and Q.ICD.39) associated with GY were identified on chromosomes 1A, 1B, 6A and 6B (Fig 4). In particular, Q.ICD.37 was also associated with GpS and SPK, while Q.ICD.39 and Q.ICD.04 also controlled TKW and GpS.

Under moisture stressed conditions, GY was associated to 14 loci (Fig 4). Among these, Q.ICD.08, Q.ICD.11, Q.ICD.17, Q.ICD.20, Q.ICD.28 and Q.ICD.44 on chromosomes 1B, 2A, 3A, 3B, 4B, 7B, respectively were identified in two or more stressed environments. Interestingly, locus Q.ICD.28 was also associated with TKW, SPK and GpS, while Q.ICD.44 controlled TKW, in addition to GY.

A comparison of significant loci for GY across stressed and non-stressed conditions identified a consistent locus on chromosome 7A (Q.ICD.41), also controlling GpS and SPK, on chr 5 A (Q.ICD.31) associated with GY, TKW, and SPK, and on chr 1A (Q.ICD.01) controlling GY, TKW, SPK and GpS. This last QTL was the most frequently identified region across all environments.

The GWAS conducted for yield stability (AWAI) revealed two QTLs (Q.ICD.02 and Q.ICD.13) on chromosomes 1A and 2B. Interestingly, both QTLs were linked to GpS and TKW. In addition, for TKW, two additional loci (Q.ICD.18 and Q.ICD.35) not associated with GY were identified on chromosomes 3A and 5B.

Conducting GWAS for the ‘confirmation panel’ confirmed the importance of Q.ICD.08, Q.ICD.11, and Q.ICD.44, which were also identified as important QTL in the ‘discovery panel’ under moisture stress-conditions, and Q.ICD.13 identified as associated to the control of yield stability (AWAI).

### Effect of different allele combination on water productivity classes

To determine the allelic effect on grain yield across environments, the main representative marker of each QTL was investigated as single marker regression at all locations (Fig 5). The major allele of AX-94549122 is strongly correlated with grain yield under both moisture conditions, while for AX-95631864 is the minor allele that is associated with both conditions. Hence, these two loci contribute to yield overall. Instead, AX-94910470 major allele is mostly important for non-stressed environments, while AX-95191125 minor allele is linked only to stressed conditions. Hence, these two loci control yield performances under different moisture conditions. The combination of major allele at all QTLs explained variation at all non-stressed environment, and only in few stressed ones.

**Figure 5.**
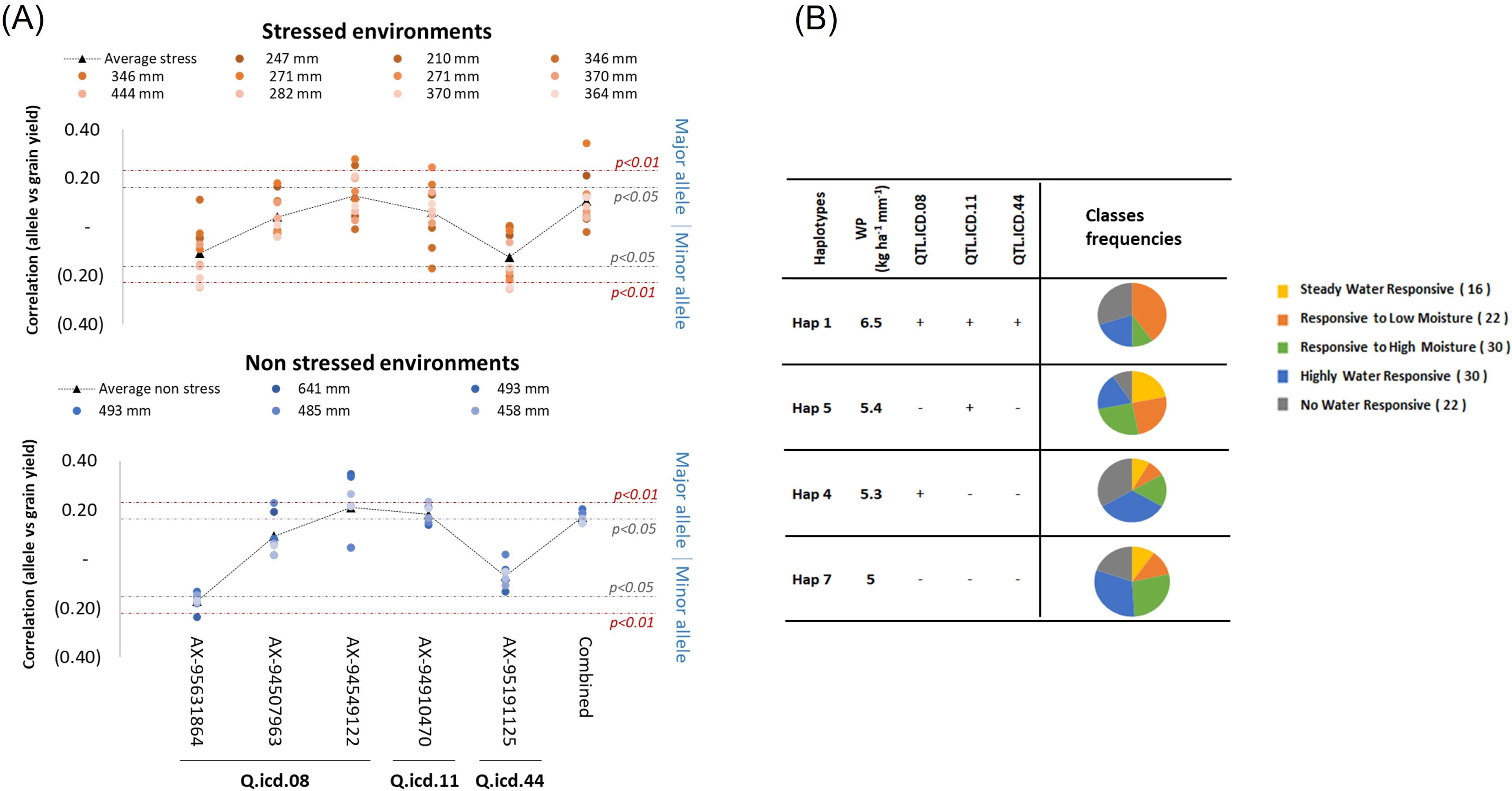
**(A).** Correlation between allelic call of five major markers representing three QTLs and grain yield at eighteen environments, presented for the moisture stressed and non-stressed separately. Each environment is named as the total of its moisture, and color coded from darker to brighter orange for severity of drought, and darker to lighter blue for decreasing moisture content. The average performance is presented as a black triangle. The Pearson’s significant cut off are presented for both major and minor alleles. **(B)**. Allelic haplotypes effects of the significant loci on water productivity of 120 genotypes grown under 18 environments. Left: The accessions were divided into four groups based on their haplotype for three major QTLs: ‘+’ mark the positive and ‘-’ the negative alleles. Right: The haplotype frequencies of each water response class.

To better assess the interaction between QTL, the ‘discovery panel’ entries assigned to the five WP classes were investigated for their haplotype composition at these three QTLs (Q.ICD.08, Q.ICD.11 and Q.ICD.44). Four haplotypes groups could be identified (Fig 5). Haplotype 1 with favorable alleles at all loci reached the highest average water productivity of 6.5 kg ha^-1^ mm^-1^, with 40% of the genotypes belonging to class 2 ‘Responsive to low moisture’. Interestingly, the same allelic combination is responsible for high grain yield under drought (Fig S3). Haplotypes 2 and 3 reach an average water productivity of respectively 5.3 and 5.4 kg ha^-1^ mm^-1^, with only one positive allele. Haplotype 2 contributes equally to the four classes. While haplotype 3 is mainly expressed by class 4 ‘highly water responsive’. Haplotype 4 harboring three negative alleles reaches 5 kg ha^-1^ mm^-1^, with classes 3 and 4 ‘responsive to high moisture’ and ‘highly water responsive’ corresponding to the highest portions of this haplotype.

### Confirmation of haplotype effect

The three main QTLs (Q.ICD.08, Q.ICD.11, and Q.ICD.44) were investigated for their additive effect within the ‘confirmation panel’. A total of five haplotypes were identified within the panel for these three loci (Fig 6). The panel was tested under moisture stress at one environment (Sidi el Aydi) during season 2018-19. The linear model confirmed that the haplotype groups represented discrete classes with significant difference. Haplotype 1 (Hap1) carrying only favorable alleles at all QTLs showed a GY advantage of more 704.6 kg ha^-1^ compared with haplotype 7 with no positive alleles at the three loci and a consequent gain in water productivity of 2.2 kg ha^-1^ mm^-1^. Also, Hap 3, with 2 positive alleles, except for Q.ICD.11 was significantly superior to Hap 7 (no positive alleles), but it was not superior to Hap 4 (only 1 positive allele). This suggests that Q.ICD.11 has the strongest effect, followed in order by 8 and 44.

**Figure 6.**
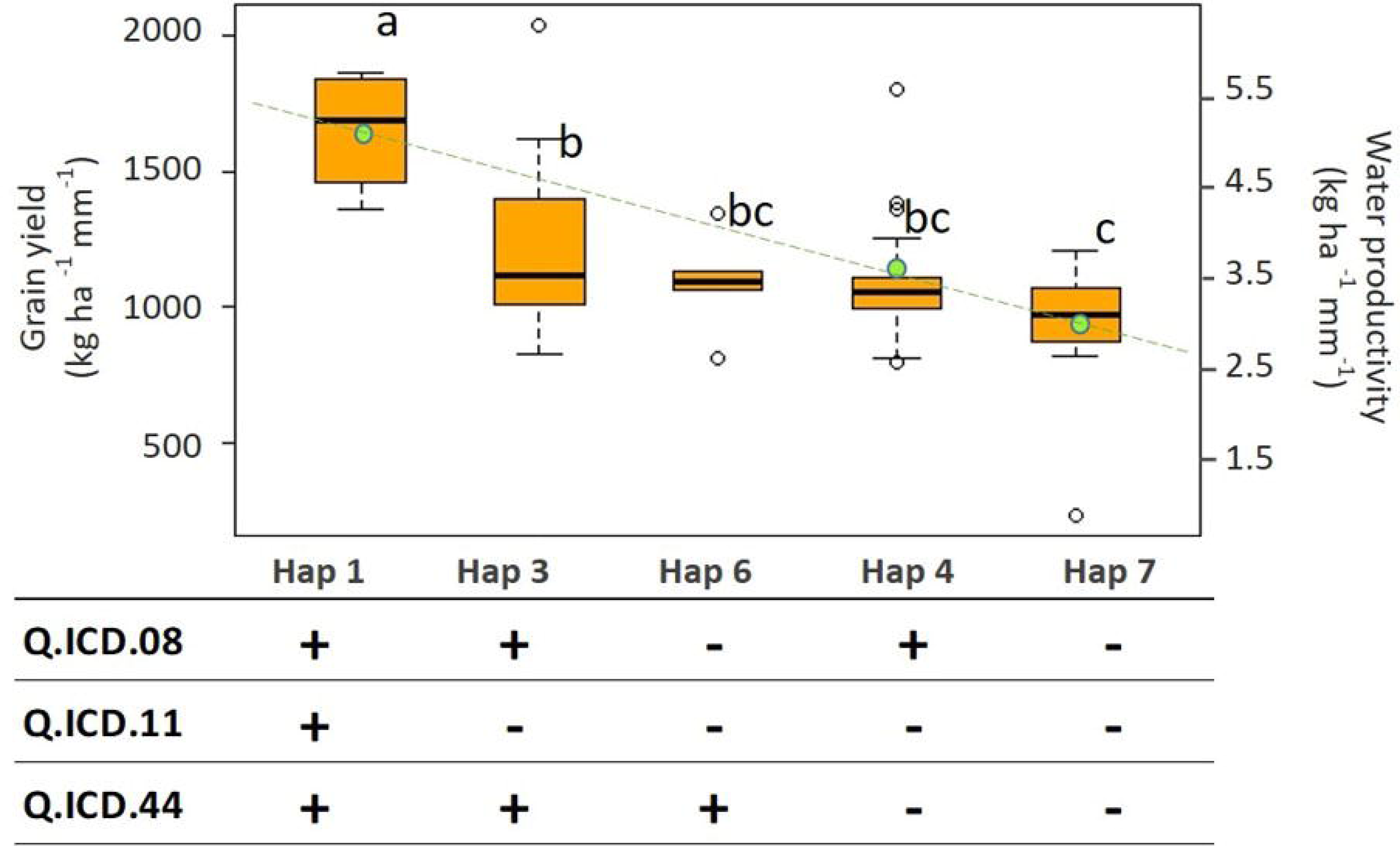
Allelic effect for the combination of 3 loci associated with GY on the ‘confirmation panel’ under moisture stress. The black line inside the boxes indicates the median of each haplotype across each cluster. The ‘+’ for the positive and ‘-’ for the negative alleles. Letters (a, b, c) indicate the LSD test. The green dots and dashed line represent the average WP.

Figure 6. Allelic effect for the combination of 3 loci associated with GY on the ‘confirmation panel’ under moisture stress. The black line inside the boxes indicates the median of each haplotype across each cluster. The ‘+’ for the positive and ‘-’ for the negative alleles. Letters (a, b, c) indicate the LSD test. The green dots and dashed line represent the average WP.

### Conversion and validation to KASP

Markers conversion and validation are quintessential steps to convert the discovery of QTLs into usable tools for breeders. Out of 36 array probes known to span the three major QTLs (Q.ICD.08, Q.ICD.11and Q.ICD.44), KASP primers could be designed for 32 of them; of these 17 were purchased and used to screen the ‘validation panel’. Nine of these detected a polymorphism within this elite set (MAF>3%). Two explained a significant (p>0.05) and three a highly significant (p<0.01) portion of the phenotypic variation for grain yield (Fig 7), when assessing the panel at the severely drought affected station of Sidi el Aydi during season 2019-20. KASP were validated for Q.ICD.08 located on chromosome 1B, and one each for Q.ICD.11 and Q.ICD.44 on chromosome 2A and 7B, respectively. All five markers are suitable for use in MAS, and their use in combination shall further increase their independent scores. In fact, AX-95631864 (Q.ICD.08) has overall the best average performance for all criteria, and the highest prediction of phenotypic variation (r^2^ = 0.10), while AX-94507963 (Q.ICD.08) is particularly suitable to identify the top yielders (true positive) with the highest overall sensitivity but it has low precision, instead AX-94549122 (Q.ICD.08) and AX-95191125 (Q.ICD.44) have perfect precision (true negative) in identifying the lines to be discarded, but low sensitivity. AX-94910470 tags the hypothetically strongest QTL (Q.ICD.11). but within the ‘validation panel’ its contribution was minor. However, the combined selection for carrying the positive allele at all five markers resulted in a drastic increase in precision, with only top performing lines being selected.

**Figure 7.**
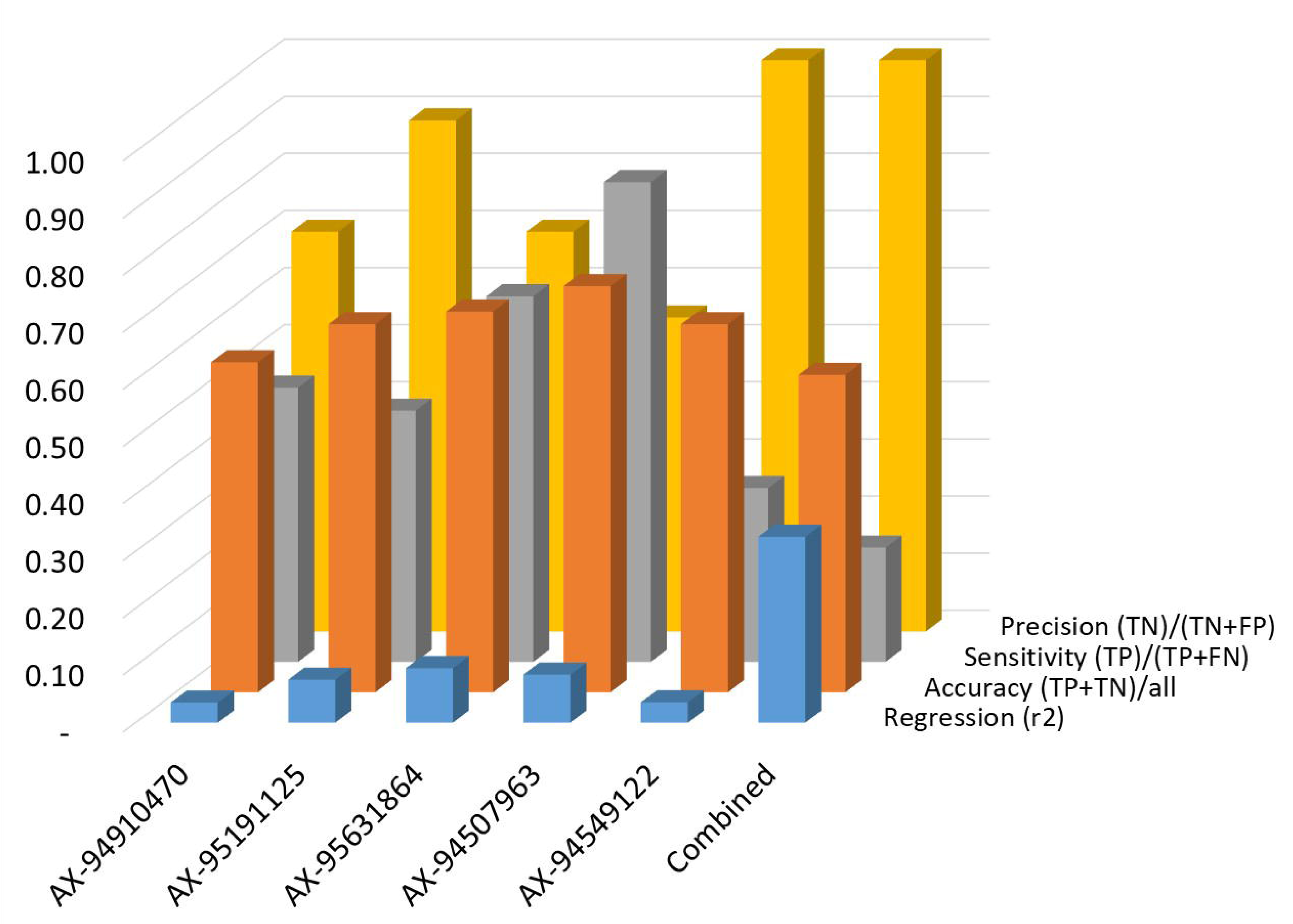
KASP markers validation on an independent set of 94 elite lines of ICARDA tested under severe drought. Correlation was measured between the BLUE for grain yield recorded at Sidi el Aydi and the haplotype score. Accuracy, sensitivity, and specificity were determined using only the top 20 and worst 20 yielding lines. AX-95631864, AX-94507963 and AX-94549122 tag Q.ICD.08, AX-94910470 tags Q.ICD.11 and AX-95191125 tags Q.ICD.44.

## Discussion

### Water productivity classes explain the genotype response to moisture stress

This study evaluated a global ‘discovery panel’ at 18 environments. Climatic regression confirmed that grain yield variation at these environments was mainly controlled by the moisture availability during the vegetative, flowering, and grain filling phases. Liliane and Charles (2020) and El Haddad *et al*. (2021) also found that moisture availability during the vegetative stage is a critical climatic factor influencing the response of durum wheat genotypes. In fact, the major negative impact of drought stress on wheat is the reduction in fresh and dry biomass production (Farooq *et al*., 2009), which affects grain number and grain size (Dickin and Wright, 2008). A common response to cope with drought stress is stomatal closure, since that also alters the photosynthetic rate, plants must constantly adjust stomatal conductance to maintain a balance between sufficient CO_2_ uptake and water loss. Toulotte *et al*. (2022) hypothesized that reduced rates of stomatal conductance and subsequently decreased water loss due to reduced stomatal density, allowed the available plant resources to be allocated to seed propagation and aboveground Biomass. A previous study (Sokoto and Singh, 2013) revealed that water stress at vegetative stage significantly reduced spike length and grains per spike. While water stress at flowering and grain filling significantly reduced 1000 kernel weight, grain yield and harvest index. In fact, water stress induced accelerated senescence after anthesis shortens the duration of grain filling by causing premature desiccation of the endosperm and by limiting embryo volume has also been reported (Westgate, 1994).

Principal component clustering based on the most critical climatic factors allowed us to classify the sites into two mega environments: moisture stressed and non-stressed. The effect of moisture was further partitioned by assigning genotypes to five classes of WP. The same set of genotypes was tested for yield potential under the two mega-environments as well as yield stability overall. Interestingly, the highly yielding genotypes under moisture stress belong to class 2 ‘Responsive to low moisture’. While under non stress, the highly yielding belongs to class 3 ‘Responsive to high moisture’. And the class 4 ‘Highly water responsive’ represent mainly the genotypes highly yielding under both conditions. Similarly, to Siahpoosh and Dehghanian (2012) genotypes were significantly different for WP. The highly phenotypic variation was due to the environmental effect. The plant response to water stress varied. The decrease of production can be due to the plant defense by reducing stomatal conductance and CO_2_ assimilation rate (Catola *et al*., 2016). While, when the plant had high water productivity under moisture shortage, Stallmann *et al*. (2020) explained this reaction by the increase of the intrinsic plant water use efficiency caused by the stomatal closure, which restrict transpiration before it inhibits photosynthesis.

Correlation analysis was done to determine the main traits contributing to grain yield under drought. Interestingly, grain yield was positively correlated with yield components in moisture stressed environments. These findings were consistent with KiliÇ and Tacettin (2010) and Al-Ghzawi *et al*. (2018), who reported that spike per m^2^, grains per spike and TKW were directly related to grain yield. Since, genetic research has shown that it is possible to increase grain size without a negative effect on grain number (Rivera-Amado *et al*., 2019). The negative correlation between yield under stress and heading date has frequently been reported (Dodig *et al*., 2010; Gonzalez-Ribot *et al*., 2017), indicating that the most precocious genotypes would be desirable in accordance with other reports for Mediterranean environments (Acevedo and Ceccarelli, 1989; Quarrie *et al*., 1999; Richards *et al*., 2001).

### Genetic dissection of drought tolerance in durum wheat

Breeding cultivars able to thrive under moisture stressed conditions is challenging, since wide adaptation is hindered by high genotype by environment interaction. Drought tolerance is a complex quantitative trait controlled by an army of loci interacting with the environment. Blanco *et al*. (2011a) reported that genomic regions linked with GY are present in all chromosomes, and that the magnitude of their effect varies based on the environment. Eleven loci were identified as responsible for the control of GY in more than one environment in our study. Four QTLs (Q.ICD.04, Q.ICD.07, Q.ICD.37, Q.ICD.39) were active under non moisture stressed conditions located on chromosomes 1A, 1B, 6A and 6B, six QTLs (Q.ICD.08, Q.ICD.11, Q.ICD.17, Q.ICD.20, Q.ICD.28, Q.ICD.44) on chromosomes 1B, 2A, 3A, 3B, 4B, 7B under moisture stress, and QTL.ICD.41 and Q.ICD.01 on chromosome 1A and 7A were common under both conditions. Sukumaran *et al*. (2018), Rahimi *et al*. (2019) and Muhu-Din Ahmed *et al*. (2020) all identified QTLs on chromosomes 1A, 6A and 6B related to GY under non moisture stressed conditions in durum wheat. Interestingly, most QTLs controlled also at least one of the yield components. In particular, Q.ICD.01 was linked to all measured four traits (TKW, SPK and GpS) and a similar region was already identified by other authors for its importance in wheat for GY under different water regimes (Charmet *et al*., 2001; Ain *et al*., 2015; Gupta *et al*., 2017; Rahimi *et al*., 2019; Muhu-Din Ahmed *et al*., 2020; Zandipour *et al*., 2020), yield stability (Sehgal *et al*., 2020), TKW (Neumann *et al*., 2010; Lozada *et al*., 2017; Ogbonnaya *et al*., 2017), GpS and SPK (El Hassouni *et al*., 2019). Similarly, to previous studies (Shokat *et al*., 2020; Xin *et al*., 2020), TKW was positively correlated with grain yield under drought and irrigated conditions, indicating that plant genotypes having higher TKW under irrigated conditions often have a chance to maintain higher TKW under drought conditions (Shokat *et al*., 2020). Less reduction in TKW will ideally allow good yield under drought conditions. Previous studies have found QTLs for TKW on almost all chromosomes of the wheat genome (McCartney *et al*., 2005; Williams *et al*., 2012; Okamoto *et al*., 2013; Zhang *et al*., 2015; Cabral *et al*., 2018; Pradhan *et al*., 2019), we found the same except on chromosome 7A. The most consistent loci have been detected on chromosomes 3A and 4B; Q.ICD.18 and Q.ICD.28 can be compared with previous finding by Pinto *et al*. (2010) and Sun *et al*. (2017). Grain per spike is correlated with grain yield under both moisture and no moisture stressed conditions (Pradhan *et al*., 2019), preserving high GpS during drought conditions is important in order to keep good yield. We found significant associations of GpS (Q.ICD.07 and Q.ICD.31) mainly with chromosomes 1B and 5A under both conditions. While significant regions for spike per m^2^ (Q.ICD.25 and Q.ICD.46) were mainly located on chromosomes 4B and 7B. For AMMI wide adaptation index, 4 MTA spanned on 2 QTLs (Q.ICD.02 and Q.ICD.13) were detected on chromosomes 1A and 2B. Contrary to the finding of Sehgal *et al*. (2017), both QTLs were not associated to GY instead were linked to TKW and grain per spike. Recently, Sehgal *et al*. (2020) identified haplotypes blocks associated with stability index Pi on chromosome 1A, using advanced bread wheat lines under contrasting environments. While the role of chromosome 2B in controlling stability was previously reported in a large elite panel of wheat (Sehgal *et al*., 2017) and a winter wheat population (Lozada and Carter, 2020), as most of the significant MTAs controlling yield trait stability were detected in this chromosomal region.

GY showed a significantly positive correlation with TKW, GpS and SPK, indicating that the increased GY under moisture stress resulted from increased yield components. While under non moisture stress, the GY increase is mainly due to the significant relationship with TKW and GpS. Consequently, it is feasible to improve GY by selecting these yield related traits in breeding programs because of the more accurate measurement across moisture and non-moisture stressed environments in comparison with yield.

The three QTLs Q.ICD.08, Q.ICD.11 and Q.ICD.44 on chromosomes 1B, 2B and 7B linked to grain yield under low moisture, were used to investigate the allelic combination responsible for water productivity. Interestingly, class 2 genotypes had mainly the positive alleles for all three loci providing a significant water productivity advantage of +1.5 kg ha^-1^ mm^-1^ under low moisture environments. While class 4 had one positive allele at AX-94910470 belonging to Q.ICD.11.

The three main QTLs were confirmed by a second independent ‘investigation panel’ grown under moisture stress. The haplotype assessment confirmed that carrying the positive alleles at all loci increased grain yield by +704.6 kg ha^-1^ and water productivity by 2.2 kg ha^-1^ mm^-1^. Juliana *et al*. (2021) by using a large scale GWAS, reported the highest number of consistent GY GBS markers on chromosomes 2A, 6B, 6A, 5B, 1B and 7B. Similarly, several studies have also reported QTLs on chromosome 1B responsible for the control of GY in durum wheat (Roncallo *et al*., 2017; Xu *et al*., 2017; Rehman Arif *et al*., 2020). The effect of Q.ICD.08 under moisture stress was also identified in Juliana *et al*. (2021) study, loci controlling GY under optimum and drought. That can be explained by Mathew *et al*. (2019) findings, who reported root and shoot biomass association region on chromosome 1B. Further, Pshenichnikova *et al*. (2021) found 53 QTLs associated with physiological and agronomic traits under contrasting water supply. Those findings may explain the importance of Q.ICD.11.

Q.ICD.44 was associated with TKW and grain yield in combined and four environments under drought represent important loci. Zaïm *et al*. (2020) found in a recent mapping populations study tested across dry environments, a consistent QTL for GY in the same chromosome. Similarly, by using a diverse population of winter wheat, Lozada and Carter (2020) found a site controlling multiple yield trait and trait stability measures in the same chromosome.

### Markers validation for marker assisted selection

Axiom to KASP marker conversion and validation was conducted for 17 MTA. Only five KASP markers generated polymorphic haplotypes in the independent set of ICARDA elites IDON43. All five demonstrated significant (p<0.05) correlation to grain yield assessed under the severely drought affected station of Sidi el Aydi. AX-95631864, AX-94507963 and AX-94549122 tag Q.ICD.08 located on chromosome 1B, one of the three main loci identified in this study. The first two markers revealed good correlation, accuracy, precision and sensitivity, while the third one has medium sensitivity. Similarly, to AX-94910470 and AX-95191125 tag respectively Q.ICD.11 and Q.ICD.44 on chromosomes 4B and 7B. Those markers are protected against Type I errors, but they are prone to Type II errors, with several lines identified as not carrying the positive allele while instead being tolerant to drought. The five markers could be considered as validated and useful for wheat breeding.

## Conclusion

Drought tolerance is a complex quantitative trait that is influenced by genetic background and highly hindered by genotype by environment interactions. To understand the mechanism and the implied loci, a panel was tested under eighteen environments, clustered as moisture stressed and no-moisture stressed environments. Our results confirmed that besides grain components, water productivity is the most critical trait to drive tolerance to moisture stress, and hence should be the primary targets of durum wheat breeders. A total of six QTLs were associated with GY under drought, some of them were linked with TKW, GpS and SPK. The haplotype diversity of three markers each from the three most promising QTLs Q.ICD.08, Q.ICD.11 and Q.ICD.44 size 19, 83 and 20 Mbp, causes water productivity of up to +1.5 kg ha^-1^ mm^-1^ across moisture stressed conditions. The three markers were validated into KASP markers and can further be utilized in marker assisted selection, beside theremaining QTLs after validation to improve the drought tolerance and yield stability in wheat. The lines Magrour (GID: 800032178), Icakassem1 (GID: 800030179) and DAWRyT-0106 (GID: 800043267) were confirmed to be drought tolerant and carrying the positive alleles of the three main QTLs. Those genotypes may serve as ideal crossing material in breeding programs.

### Supplementary data

Figure S1. Genome-wide average linkage disequilibrium (LD) decay over genetic distances of the second discovery set. Plot of pair-wise LD r^2^ values as a function of inter-marker map distance (Mbp). The blue curve represents the model fit to LD decay. The red line represents the intercept to r^2^ =LJ0.2. a) for the discovery panel. b) for the investigation set.

Figure S2. Boxplots of grain yield (GY) performances and its components across environments. The medians are indicated by black line inside the boxes. The box borders indicate upper and lower quartiles, the caps indicate 90^th^ and 10^th^ percentiles, and the circles indicate observations below and above those percentiles. TKW: 1,000 kernel weight, SPK: spike density per m^2^, GS: grain per spike.

Figure S3. Average response of durum wheat water productivity across different rates of moisture.

Figure S4.LJAllelic effect for the combination of the 3 loci associated with GY under moisture stress.

Table S1. List of the “discovery” panel and its kinship assignment (k=10)

Table S2. List of the “investigation” panel IDON42 and its kinship assignment (k=8)

Table S3. List of the “validation” panel IDON43.

Table S4. Analysis of variance for GY, DTH, TKW, SPK, GpS in 18 environments.

Table S5. Linear regression between grain yield and climatic factors across eighteen environments and four five seasonal timepoints.

Table S6. Moisture classes assignments of the 121 selected genotypes and their water productivity characteristics under stressed and no-stressed conditions.

Table S7. Markers associated with the tested traits under no- and moisture stressed mega-environments.

## Acknowledgments

We wish to acknowledge Rached Abdelaziz and the other field technical staff in Morocco, Lebanon and Jordan for the intense support.

## Author contributions

Conceptualization and methodology, M.Z. and F.M.B.; Statistical analysis, M.Z. and Z.K.; validation, F.M.B.; writing original draft preparation, M.Z.; project administration, F.M.B. All authors have read and agreed to the published version of the manuscript.

## Data availability

The germplasm described here is available through ICARDA’s genebank and can be requested here: https://www.genesys-pgr.org/wiews/SYR002. The genotypic and phenotypic data have been provided as supplementary.

## Abbreviations

AMMI: additive main effect and multiplicative interaction
AWAI: AMMI wide adaptation index
bi: slope value
BLUE: Best linear unbiased estimates
DTH: Days to heading
E: environment
G: genotype
GxE: genotype by environment
GpS: Grain per spike, interaction
GWAS: genome wide association study
GY: grain yield
IDON: International Durum Observatory Nursery
KASP: Kompetitive Allele Specific PCR
KFD: Kfardan
MTA: marker trait association
MCH: Marchouch
MKZ: Melk Zhar
MUS: Musghar
PCA: principal component differentiation
PCR: Polymerase chain reaction
PLH: Plant height
SPK: Spike density per m^2^
QTL: quantitative trait loci
RGA: root growth angle
SAD: Sidi El Aidi
TER: Terbol
TES: Tessaout
TKW: 1,000 Kernel weight
WP: water productivity
Z: Zadok’s scale.

